# In-depth and 3-Dimensional Exploration of the Budding Yeast Phosphoproteome

**DOI:** 10.1101/700070

**Authors:** Michael Charles Lanz, Kumar Yugandhar, Shagun Gupta, Ethan Sanford, Vitor Faça, Stephanie Vega, Aaron Joiner, Chris Fromme, Haiyuan Yu, Marcus Bustamante Smolka

**Affiliations:** Department of Molecular Biology and Genetics, Weill Institute for Cell and Molecular Biology, Cornell University, Ithaca, NY 14853; Department of Computational Biology, Weill Institute for Cell and Molecular Biology, Cornell University, Ithaca, NY 14853

## Abstract

Phosphorylation is one of the most dynamic and widespread post-translational modifications regulating virtually every aspect of eukaryotic cell biology. Here we present a comprehensive phosphoproteomic dataset for budding yeast, comprised of over 30,000 high confidence phosphorylation sites identified by mass spectrometry. This single dataset nearly doubles the size of the known phosphoproteome in budding yeast and defines a set of cell cycle-regulated phosphorylation events. With the goal of enhancing the identification of functional phosphorylation events, we performed computational positioning of phosphorylation sites on available 3D protein structures and systematically identified events predicted to regulate protein complex architecture. Results reveal a large number of phosphorylation sites mapping to or near protein interaction interfaces, many of which result in steric or electrostatic “clashes” predicted to disrupt the interaction. Phosphorylation site mutants experimentally validate our predictions and support a role for phosphorylation in negatively regulating protein-protein interactions. With the advancement of Cryo-EM and the increasing number of available structures, our approach should help drive the functional and spatial exploration of the phosphoproteome.

## Introduction

Post-translational modification of proteins by phosphorylation controls virtually every cellular process. Regulatory mechanisms based on phosphorylation have been widely explored and characterized. In classical approaches, phosphorylation sites are often biochemically identified on substrate proteins of interest and then mutated to either prevent or constitutively mimic a phosphorylation event in order to determine the biological relevance. The phenotypes associated with these “phosphomutant” proteins inform on the biological purpose of phosphorylation at that site. In the last 15 years, advances in mass spectrometry have greatly expanded our ability to identify phosphorylation events, leading to large phosphoproteomic databases ^1–5^. However, our ability to probe the biological relevance of the identified phosphorylation events still relies on low throughput methods. As a consequence, the functional importance of most catalogued phosphorylation events has not yet been determined ^6^. Notably, given the overwhelming number of identified phosphorylation events, over 100,000 in the case of a human cell (PhosphoSitePlus ^7^), and likely promiscuity in kinase actions, it is debatable whether all of these events are functionally relevant ^8, 9^. In many cases where attempts have been made to investigate the role of specific phosphorylation events, the results are often negative ^10^, consistent with the notion that many phosphorylation events may be extensively redundant in nature or, perhaps, not functional ^9, 11^. These issues highlight the necessity for strategies to predict functional phosphorylation sites from large phosphoproteome datasets. While guidelines for interpreting phosphoproteomic data sets to identify candidate sites for mutational analysis are available ^10^, strategies to efficiently and systematically identify functional phosphorylation events are lacking.

Here we present an in-depth phosphoproteome for budding yeast that constitutes, to the best of our knowledge, the single largest collection of phosphorylation sites for this organism. Over 10.6 million high resolution MS/MS spectra were acquired in our mass spectrometer and subjected to a parallel search approach to maximize the number of phosphosites identified with high-confidence. In addition, we utilized two independent methods for scoring phosphosite localization and employed an in-house algorithm to capture ambiguous phosphosites that fall within clusters of consecutive, phosphorylate-able residues. Remarkably, our dataset nearly doubles the size of the budding yeast phosphoproteome. With the goal of improving the systematic prediction of functional phosphorylation events, we computationally positioned phosphorylation onto all available 3D protein structures and systematically identified potentially functional phosphorylation events. Results reveal a large number of phosphorylation sites mapping to or near protein interaction interfaces, some of which result in steric or electrostatic “clashes” predicted to disrupt the interaction. Phosphorylation site mutants experimentally validate our predictions and establish roles for phosphorylation in negatively regulating protein-protein interactions. We have compiled our in-depth phosphoproteome into an on-line database open to the community. This resource should help drive the functional and spatial exploration of the phosphoproteome.

## Results

### In depth phosphoproteome of budding yeast

We sought to generate an in-depth phosphoproteomic database for the model system budding yeast. Since most currently available phosphosite databases consist of data deposited from multiple independent groups, each utilizing their own instrumentation and acquisition / spectral search methods, we aimed to generate a repository of phosphorylation events sourced strictly from high-resolution spectral data acquired using a single, in-house mass spectrometer and processed through a unified data processing pipeline (Fig. 1). The spectral set used to assemble the database was generated from 75 independent SILAC-based experiments, all of which consisted of IMAC-mediated enrichment of phosphopeptides from whole cell lysates fractionated by HILIC chromatography (Fig. 1). The experiments that contributed to our dataset were originally performed for various unrelated biological inquiries, and explored a range of conditions, including distinct cell cycle stages, DNA damage treatment and carbon deprivation (Fig. 1). A fraction of this dataset was previously published ^12^. In all, the spectral library consisted of fragmentation spectra acquired from over 825 independent MS runs (1500+ hours of data-dependent acquisition time). To identify Peptide Spectrum Matches (PSMs) from our spectral library, we utilized three different search engines (SORCERER, Proteome Discoverer (PD), and MaxQuant (MQ)) (Fig. 1; see details under Materials and Methods). A critical challenge in the analysis of peptide-centric phosphoproteomic experiments is the need to properly assign the phosphorylated STY residue within a fragmented peptide ^13^, thus the use of multiple search engines allowed us to employ two prominent algorithms for determining phosphosite localization and maximized our ability to localize phosphorylated residues with high confidence. Additionally, we found our Sequest search engines to be more amenable for searching a massive spectral set using semi-specific tryptic digestion parameters (Fig. 1). The resulting database contains over 30,000 phosphorylation sites identified with high-confidence site localization and represents the largest available phosphoproteome dataset for this organism (Supplemental Table 1). In addition to the 30,906 sites identified with high confidence localization, we used an in-house clustering algorithm to capture several thousand more “phosphosites” whose site localization scores were distributed within consecutive STY residues (Supplemental Figure 1 highlights the rationale behind our clustering algorithm). When also considering these ambiguous phosphosites that are located within clusters of consecutive phosphorylatable residues, the total number of phosphosites identified in our dataset exceeds 36,000 (Supplemental Table 1). We acknowledge that the number of phosphosites added from the clustering step might be inflated, since in some cases multiple ambiguous phosphosites are represented within the same cluster of residues (Supplemental Table 1; see phosphosites assigned to Gic1-T221 and Gic1-S222 as an example). Consistent with the stochastic nature of data-dependent LC-MS/MS acquisition, a significant fraction of phosphosites from our study were identified from only a single phosphopeptide identification (Figure 1, Pie chart). For sites with a just single ID, we advise that the quality metric provided in Supplemental Table 1 (“Score” / “Xcorr”) be carefully considered.

**Figure 1:**
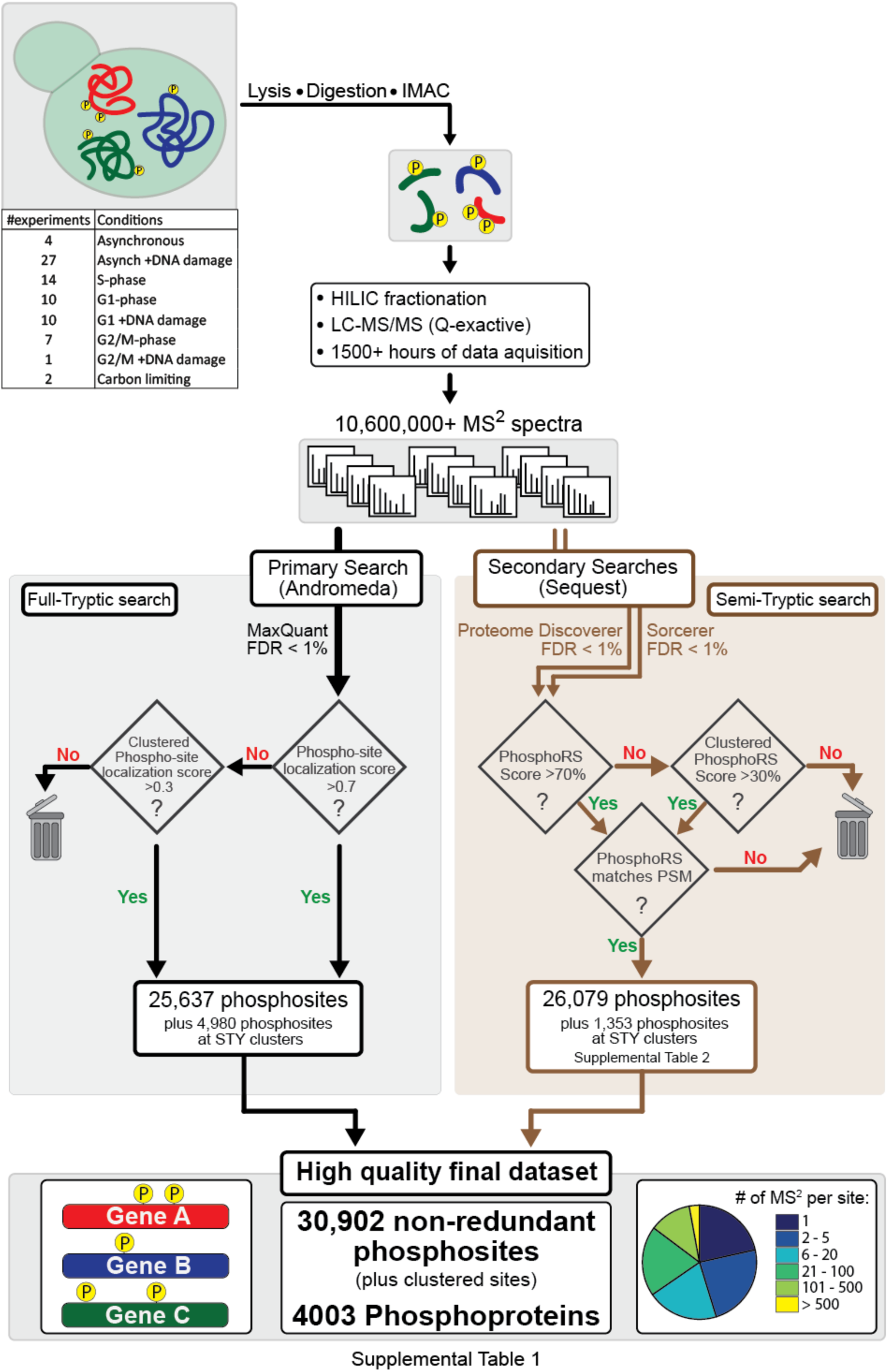
An in-depth phosphoproteome database for budding yeast. A general workflow for mapping the budding yeast phosphoproteome using mass spectrometry. The phosphoproteomic dataset was generated from a multitude of different experimental conditions, including distinct cell cycle stages. Phosphopeptides were enriched directly from trypsin- or chymotrypsin-digested cell lysates via immobilized metal ion chromatography (IMAC). All samples were highly enriched for phosphorylated peptides (80-95% phosphopeptides) and, in most cases, extensively pre-fractionated via HILIC. Phosphopeptide fragments were captured as high resolution MS^2^ spectra using an obitrap mass analyzer (Q-exactive). Three separate search engines were used to identify phosphopeptides from the fragmentation spectra. The primary search was performed using MaxQuant (Andromeda engine). Phosphosites from the primary search were extracted from MaxQuants’s “Phospho STY” output table. A secondary search was performed using two separate Sequest-based engines, Proteome Discoverer (PD) and SORCERER. The secondary search utilized similar search parameters as the primary search, with the exception that tryptic enzyme digestion was set to semi-specific. To further increase the confidence in our Sequest searches, we only considered phosphopeptides whose backbone sequence appeared in both the PD and SORCERER searches. Phosphosite localization probabilities were determined using MaxQuant (localization score) and the PhosphoRS node within Proteome Discoverer. Phosphosites with localization scores / phosphoRS scores above 70% were considered to have “high-confidence localization.” A clustering algorithm was used to capture addition phosphosites that meet our requirement for high-confidence localization (see Supplement Figure 1 for a demonstration of the logic used to cluster phosphosites). Because PhosphoRS is run independently of the PD PSM search, its phosphosite localization can sometimes conflict with the localization assigned in the rank 1 PSM. To further ensure data quality, we required agreement between the localizations determined by PD’s PSM search and PhosphoRS node. PD and SORCERER searches are considered secondary because their PSM output is only included in the final data set (Supplemental Table_1) if a phosphosite was not already identified in the primary search.

We next sought to determine the extent to which our dataset overlaps with, and expands, the previously available budding yeast phosphoproteome. We first defined what collection of phosphorylation events comprised the “previously known” phosphoproteome. In recent years, the primary repository for experimentally verified phosphorylation sites in budding yeast has been BioGRID, which contains nearly 20,000 annotated phosphosites (previously known as PhosphoGRID) ^14^. BioGRID is comprised of phosphosites identified from high throughput MS-based studies (similar to this study) in addition to phosphosites identified from low throughput investigations of individual proteins or protein complexes. However, BioGRID does not contain phosphosites identified by a more recent phosphoproteomic screen performed by Swaney et al.^4^, which, prior to this study, was the largest single phosphoproteomic dataset generated for budding yeast. When added to the BioGRID compendium, Swaney et al. contributed over 3,000 unique phosphosites (Supplemental Fig. 2). We consider the phosphosites contained within BioGRID and Swaney et al. to represent what was the “previously known” budding yeast phosphoproteome (referred from here on as BioGRID/Swaney), as together they account for nearly every phosphorylation site identification reported in budding yeast prior to our study.

To assess how the coverage of our dataset compared to what was the known budding yeast phosphoproteome, we overlaid the phosphosites identified in our study with those contained in BioGRID/Swaney (Fig. 2a,b). Strikingly, in addition to capturing almost 2/3 of the sites contained within BioGRID/Swaney (Fig. 2b), our study also identified over 17,000 novel phosphosites with high-confidence site localization (Fig. 2a). If the sites captured by our clustering algorithm are considered in addition to the 17,000 sites with high confidence localization (Fig. 2b), this study nearly doubles the size of the budding yeast phosphoproteome.

**Figure 2:**
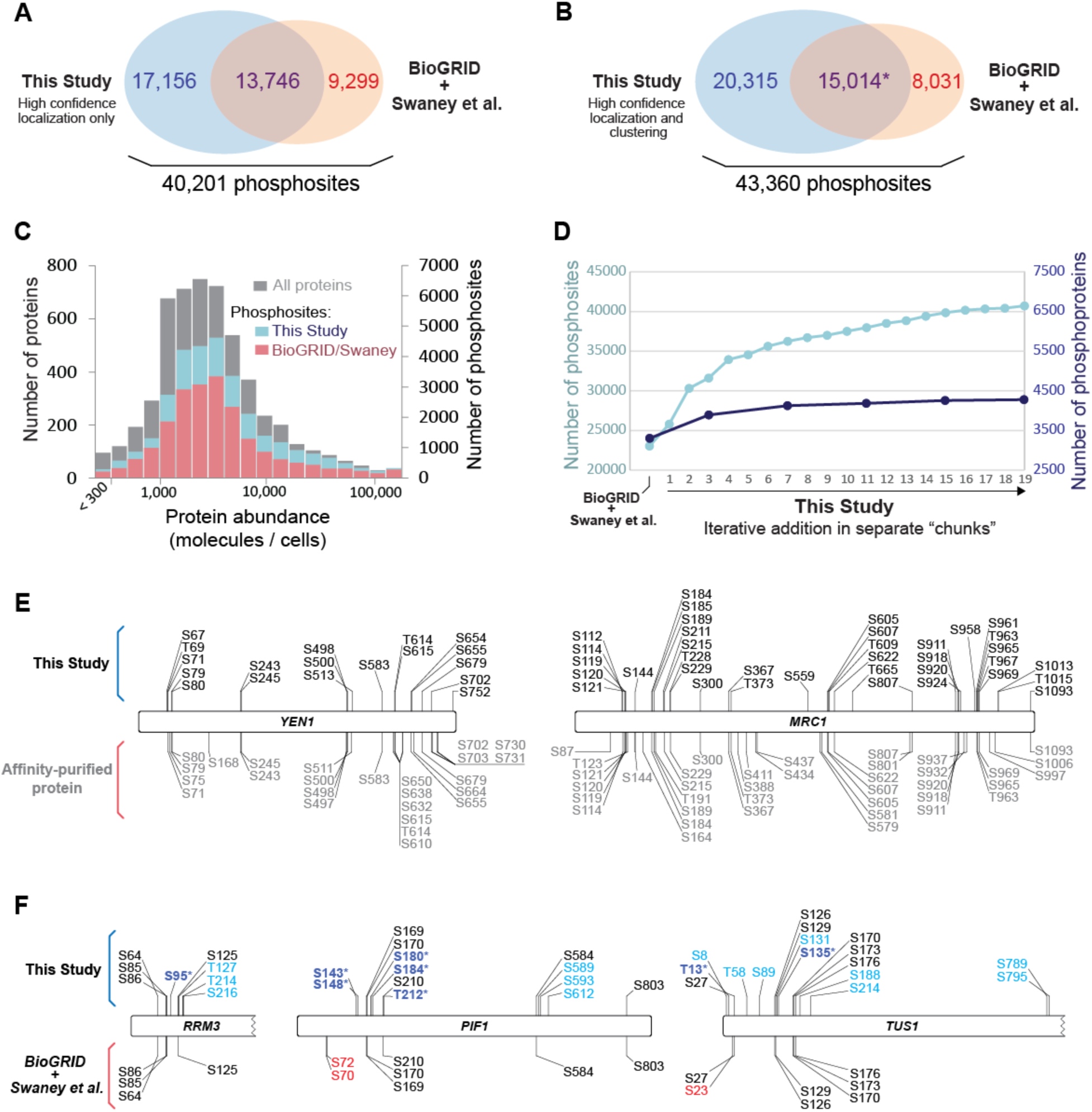
Comparative and depth analysis of current and previous budding yeast phosphoproteome datasets. (A) Venn diagrams depicting the overlap of unique phosphosites contained within our dataset with the “known” phosphoproteome prior to this study. For our dataset, only non-redundant sites (i.e. sites identified from phosphopeptides that map uniquely to a single protein) with high-confidence localization are considered. (B) As is (A) except with the inclusion of the sites captured by our clustering algorithm (Supplemental Figure 1). In this comparison, sites that redundantly mapped to multiple proteins were considered *only* when identifying overlap with the BioGRID/Swaney dataset (denoted by the asterisk) and were not considered in the 20,315 sites discovered by this study. (C) Histogram depicting the distribution of identified phosphosites as a function of protein copy number. Bars representing the number of phosphosites identified in this study are plotted behind (not on top of) the bars representing BioGRID/Swaney. (D) Dot graph assessing saturation in the ability to identify novel phosphoproteins and phosphosites from the budding yeast phosphoproteome. Unique, non-redundant phosphosites from this study were iteratively added to the BioGRID/Swaney compendium (left to right) in randomized chunks. (E) Coverage maps comparing the Yen1 and Mrc1 phosphosites identified in this study (above, in black) with the sites identified in low-throughput studies (below, in gray). For the low-throughput MS analyses, phosphopeptides were enriched after affinity purification of Yen1 (et al.) or Mrc1 (et al.) from yeast lysates. (F) Coverage maps comparing the phosphosites identified in this study (above, exclusive to this study in blue) with those found within BioGRID/Swaney (below, exclusive to BioGRID/Swaney in red). The bolded blue sites denoted by an asterisk represent putative phosphosites that were mutated an analyzed in previous studies (but were not included in BioGRID).

To further characterize the phosphorylation events being revealed by our study, we plotted our identified phosphosites as a function of protein abundance (Fig. 2c) ^15^. Despite the fact that the enrichment of phosphopeptides directly from cell lysates can hinder the detection of phosphorylation events that occur in low abundant proteins ^16^, we readily identified novel phosphorylation events in very low abundant yeast proteins, and the distribution of phosphosite discovery was mostly independent of the estimated protein abundance (Fig. 2c).

With the addition of such a large inventory of new phosphosites to an already large database, we sought to estimate whether our ability to discover new phosphoproteins or phosphosites in budding yeast is reaching saturation. Similar to what has been done previously ^17^, and using the BioGRID/Swaney dataset as a foundation, we iteratively incorporated our dataset in a randomized, “chunk”-wise manner and observed that, as the final portions of our dataset were considered, the ability to detect new phosphoproteins and phosphosites was approaching saturation (Fig. 2d). We note that the discovery of novel phosphoproteins in this study was also likely impacted by more recent updates to the annotation of the budding yeast genome/proteome. Interestingly, Albuquerque et al. previously demonstrated that the extent of phosphorylation in identified high-throughput studies is significantly less than that which can be detected from affinity purified proteins ^18^. We reasoned that if our ability to discover phosphorylation was truly approaching saturation, we should be achieving coverage at depth comparable to low-throughput MS analyses on affinity purified proteins. One such low-throughput study identified 25 phosphosites in Yen1 (Fig. 2e) ^19^, a nuclease regulated by cyclin-dependent kinase. Remarkably, we were able to detect 18 phosphosites in Yen1 (Fig. 2e), nearly all of which were identified by Blanco and colleagues. The sites identified by Blanco et al. are not currently part of BioGRID, and BioGRID/Swaney contained only four Yen1 phosphosites. Another study identified 39 phosphosites in the replisome protein Mrc1 ^18^, a number comparable to the 36 sites identified in our study (Fig. 2f). Together, these examples illustrate that, in some cases, our depth of coverage compares to depth achieved in the analysis of affinity purified proteins. In addition, we note that the depth achieved by our analysis confirmed, for the first time, the presence of phosphorylation at putative phosphosites, whose mutation was previously shown to preclude phosphorylation-dependent mobility shifts and disrupt genuine phospho-mediated regulation (Fig. 2h, bolded dark blue sites with asterisk) ^20, 21^.

### Functional and regulatory exploration of the budding yeast phosphoproteome

Despite the sheer quantity of phosphorylation revealed by mass spectrometry, the inability to distinguish meaningful phosphorylation events from “noise” within the phosphoproteome represents a fundamental limitation of the technology. To address this limitation, we employed a variety of strategies to systematically reveal potentially meaningful phosphorylation events. First, we took advantage of an extensive compilation of temperature sensitive (ts) budding yeast mutants ^22^. We reasoned that, since ts mutations fall within chemically sensitive regions of a protein’s structure, phosphorylation events which occur at or near these ts residues are more likely to be impactful. Our analysis revealed dozens of phosphorylation events that occur in immediate proximity to residues that harbor ts mutations (Supplemental Table 3), and in several cases the ts residue is itself phosphorylated. In one such case, *rsp5-T104A*, the sensitizing mutation is the substitution of a phosphorylatable threonine to alanine (Supplemental Table 3), which suggests that the phosphorylation of the Rsp5 ubiquitin ligase at T104 is somehow critical for its function.

Because phosphorylation that is subjected to dynamic regulation is more likely to be functionally important ^23^, we next aimed to extract regulatory information for the phosphorylation events we identified. While the experiments that comprise this resource were not originally designed to precisely and systematically monitor phosphorylation dynamics across the cell cycle, we were still able to obtain cycle-related regulatory information for nearly 11,000 phosphorylation events using a curation of experiments from our dataset. We compared the relative prevalence of phosphopeptides identified from yeast that were synchronized within 3 distinct cell-cycle stages (Fig. 3a). In doing so, we were able to identify thousands of phosphorylation events whose prevalence fluctuates during the cell cycle. Approximately 20 % of the phosphorylation events were either significantly enriched or depleted in one particular stage of the cell cycle (Fig. 3b; Supplemental Table 4), a proportion similar to what has been observed in fission yeast phosphoproteome ^24^. Many phosphorylation events that were subjected to cell-cycle regulation were either established cell stage-specific events or occurred within proteins with cell cycle-related functions (highlighted in Fig. 3c). Despite having less temporal resolution than more focused investigations of mitotic-specific phosphorylation dynamics ^25^, to our knowledge this analysis currently represents the most extensive catalog of cell cycle-dependent phosphorylation events for this organism at various cell cycle stages.

**Figure 3:**
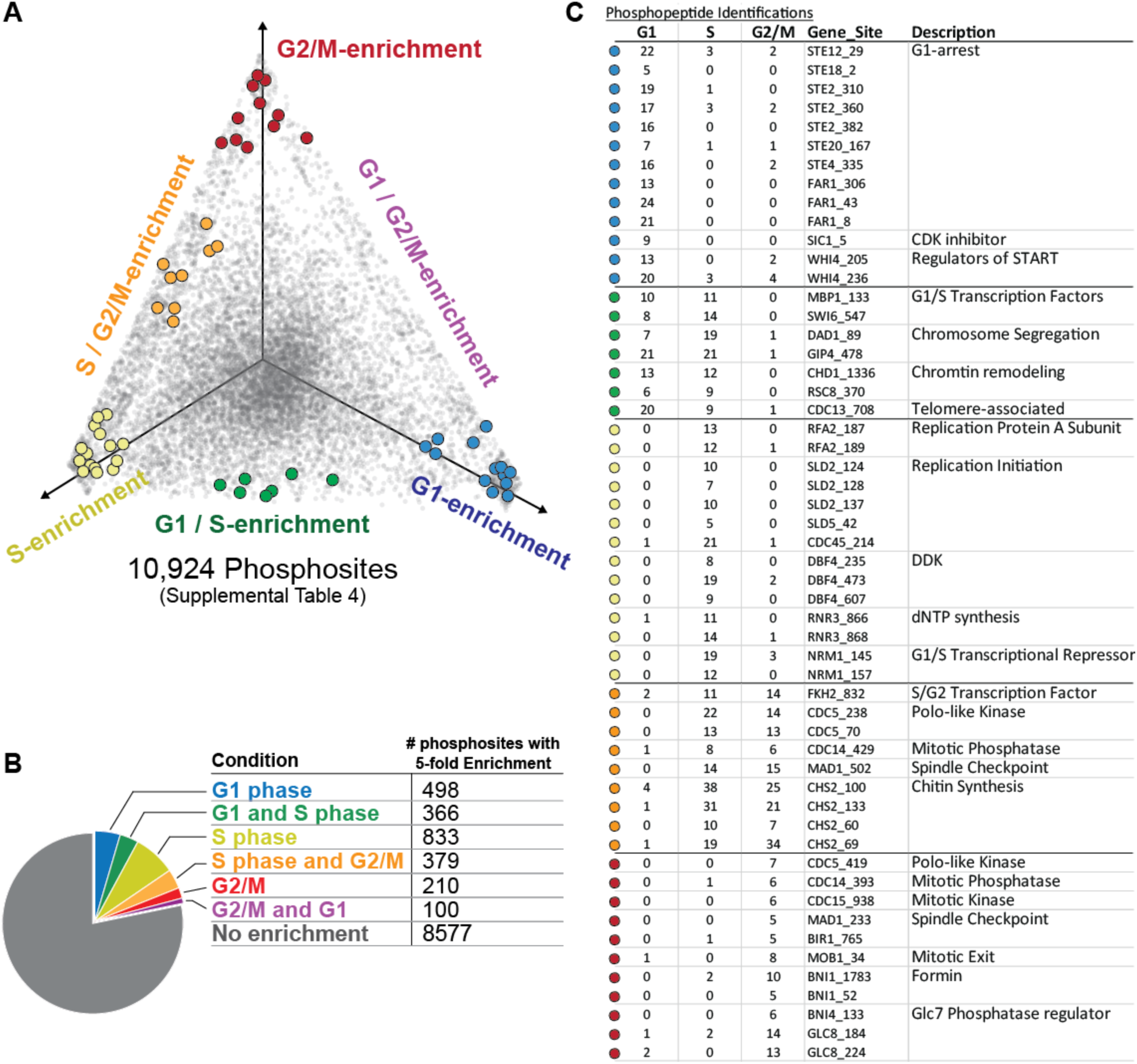
Cell cycle dynamics of the budding yeast phosphoproteome. (A) Ternary plot displaying the distribution phosphorylation events as a function of their detection in different stages of the cell cycle. A curated set of experiments from the larger dataset presented in Figure 2 was used for this analysis. Each gray dot represents a unique phosphosite (considering only the most prevalent phosphopeptide). The position of each dot within the plot represents the fraction of times it was detected in either G1, S phase, or G2/M. An 8% jitter was added to help visualize overlapping data points. (B) Table that corresponds to the highlighted dots from (A) and the number of times they were detected in each cell cycle stage. See Supplemental Table 4 for the full dataset. (C) Pie chart illustrating the fraction of phosphosites with enrichment or depletion in a particular cell cycle stage. To be considered enriched or depleted, a phosphopeptide must have 5-fold more or less detections in one particular cell cycle stage VS the other two stages (e.g. “S phase and G2/M” could alternatively be considered as “G1-depleted”). The examples given in (B) all fit the criteria for enrichment or depletion.

### The 3D budding yeast phosphoproteome

In addition to probing the regulation of phosphorylation, we performed structural analysis of phosphorylation site position with the goal of improving the systematic prediction of functional phosphorylation events. Because phosphorylation events occurring at or near protein-protein interfaces may also be more likely to impact protein function, we first plotted the phosphoproteome within the context of protein-protein interactions. We utilized Interactome INSIDER ^26^, a tool that has previously been used to systematically identify, proteome-wide, every amino acid residue found at the interface between proteins in a crystal structure or homology model (interactome3D is the source for our homology models ^27^). In total, we identified 646 phosphosites that lie at the surface between interacting proteins and many more sites that fall within 5 a.a. residues of an interface (Supplemental Table 5). An additional feature of Interactome INSIDER is its ability to predict interface residues for known protein-protein interactions that do not currently have any structural information ^26^; we found 1932 phosphorylation events that occur on these predicted protein-protein interface residues (Supplemental Table 5). This information has been compiled into the SuperPhos database (under development), which lists all identified phosphorylation sites in budding yeast and displays their proximity to known, or predicted, protein interaction interfaces (Fig. 4a).

**Figure 4:**
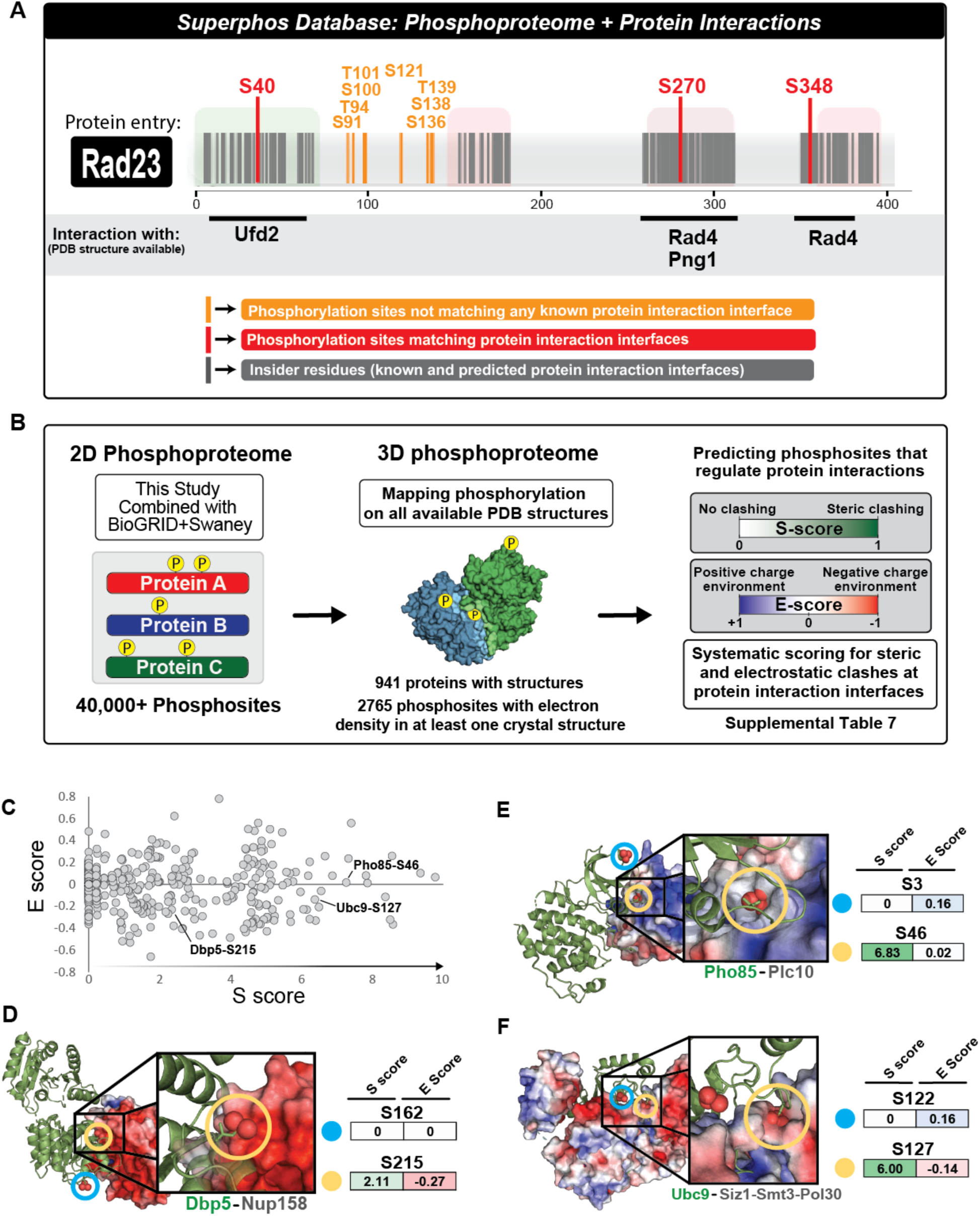
3D analysis of budding yeast phosphoproteome reveals potential regulatory phosphorylation at protein interaction interfaces. (A) The SuperPhos database (under development). Representative example of data display for a protein entry. The database merges an updated version of the budding yeast phosphoproteome with the protein interface calculations made by interactome INSIDER. In addition to indicating interactions for which PDB structures are available (as shown in this example), the SuperPhos database also provides information on all predicted interaction interfaces from INSIDER (not shown in this example). (B) Mapping the yeast phosphoproteome to all available PDB structures and systematic prediction of phosphorylation that regulates protein-protein interactions. For each phosphosite that mapped to a structured region, E and S scores were systematically calculated based on the proximity and charge of atoms from neighboring proteins. See methods for a detailed explanation of how the scores were calculated. The dot plot depicts the distribution of S and E scores assigned to each phosphosites. Phosphosites that map to more than one crystal structure or to multiple chains within a single crystal structure were assigned multiple S and E scores (Supplemental Table 7). (C) Distribution of S and E scores assigned to phosphosites in the 3D budding yeast phosphoproteome. (D), (E), (F) Representative examples of the mapping and scoring of phosphosites within the structural context of protein complexes. The phosphoprotein is displayed as a green ribbon cartoon; the electron density of the surrounding protein(s) is colored based on the electrostatic environment (as calculated by ABPS).

To further exploit available structural information in our effort to systematically identify functional phosphorylation sites, we computationally positioned the budding yeast phosphoproteome onto all available 3D protein structures within the Protein Data Bank (PDB) (Fig. 4b). When considering 941 yeast proteins with structural information in PDB with resolution better than 4 Å, we found that the majority of phosphorylation occurs only within in regions with no structural information (5,943 of 8,708 phosphosites), a finding consistent with the importance of intrinsic disorder for protein phosphorylation ^28^. Despite this, we were still able to map 2765 phosphorylation events onto structured regions (Fig. 4b). We reasoned that, because most crystallographic structures are prepared under conditions in which the crystalized proteins would not be phosphorylated (e.g. protein purified from bacteria), phosphorylation sites from our database that map to solvent-inaccessible regions within these protein structures would have a higher likely-hood of being biologically impactful. We distinguished between two types of solvent inaccessible residues: 1) residues buried within the core of a single polypeptide chain and 2) residues that lie at the interface between interacting proteins (similar to our INSIDER approach). We identified 539 phosphorylation sites that mapped to a buried, solvent-inaccessible region within a single protein (Supplemental Table 6). However, we opted to focus more on phosphorylation sites found at the interfaces where proteins interact, since these sites potentially play key regulatory roles.

### Mapping phosphorylation sites to protein interaction interfaces reveals phosphorylation events that regulate protein-protein interactions

Due to the potentially disruptive nature of adding a bulky and negatively charged phosphate group to an S/T/Y residue near a protein interaction surface, the presence of phosphorylation at a protein-protein interface could result in a steric or electrostatic clash. In these instances, we predict that interface phosphorylation would disrupt or prevent protein-protein interactions and therefore reflect a potentially important regulatory mechanism with biological implications. To systematically identify phosphorylation that would result in “clashes” between interacting proteins, we devised a minimal scoring system based on the steric and electrostatic environment surrounding phosphosites near a protein interface region (see methods for detailed explanation of how the scores were calculated). In brief, our method utilizes the per-atom charge calculated by employing PDB2PQR pipeline ^29^. Here, steric clash (S) and electrostatic (E) scores for a given phosphosite are calculated based on the distance and charge of atoms from neighboring proteins (Fig. 4c, Methods). Upon manual inspection of several phosphorylation events within their 3D context, we found that a phosphosite’s S and E scores accurately represent the surrounding steric and electrostatic environment (Figs. 4d, c, and f). We caution that the quality of our predictions is dependent on the quality and content of the structural information deposited on PDB, which can vary from structure to structure.

Using our scoring system, we extracted from our dataset hundreds of sites that, if phosphorylated in the context of the crystal structure, would cause steric clashing, occur within a negatively charged environment, or both (Fig. 4c). We hypothesized that phosphorylation events with high S scores and low E scores may disrupt protein-protein interactions. To test this hypothesis, we constitutively mimicked phosphorylation by mutating a phosphosite residue from serine to aspartic acid and determined how that mutation impacted a predicted set of protein-protein interactions. We performed proof-of-principle experiments in Rad23, an evolutionarily conserved protein with dual roles in nucleotide excision repair and proteolysis ^30^. For its DNA repair functions, Rad23, together with Rad4, recognizes damaged DNA ^31^. For its role in protein degradation, Rad23 interacts with several proteins in the ubiquitin pathway ^30^. One of these proteins is Png1, a protein involved in the degradation of misfolded ER proteins ^32^. We chose Rad23 for our proof-of-principle experiments because it harbored a phosphosite (serine 270; S270) that mapped to an interface that binds two different proteins, Rad4 and Png1 (Fig. 5a). Based on its S and E scores, we anticipated that phosphorylation at S270 might significantly impact Rad23’s interaction with Png1 while having a milder effect on its interaction with Rad4. To test our prediction, we generated a phospho-mimetic RAD23 mutant (Rad23^S270D^) and performed IP-MS to quantitatively compare interacting proteins pulled down with Rad23^WT^ vs Rad23^S270D^. Consistent with our prediction, we found that Rad23’s interaction with Png1 was specifically disrupted when phosphorylation was mimicked at S270 (greater that 25-fold change in the average SILAC ratios for Png1 peptides) (Figs. 5b,c). Importantly, the ability of Rad23 to bind to its other interacting proteins, including Rad4, was not disrupted by the mutation of serine 270.

**Figure 5:**
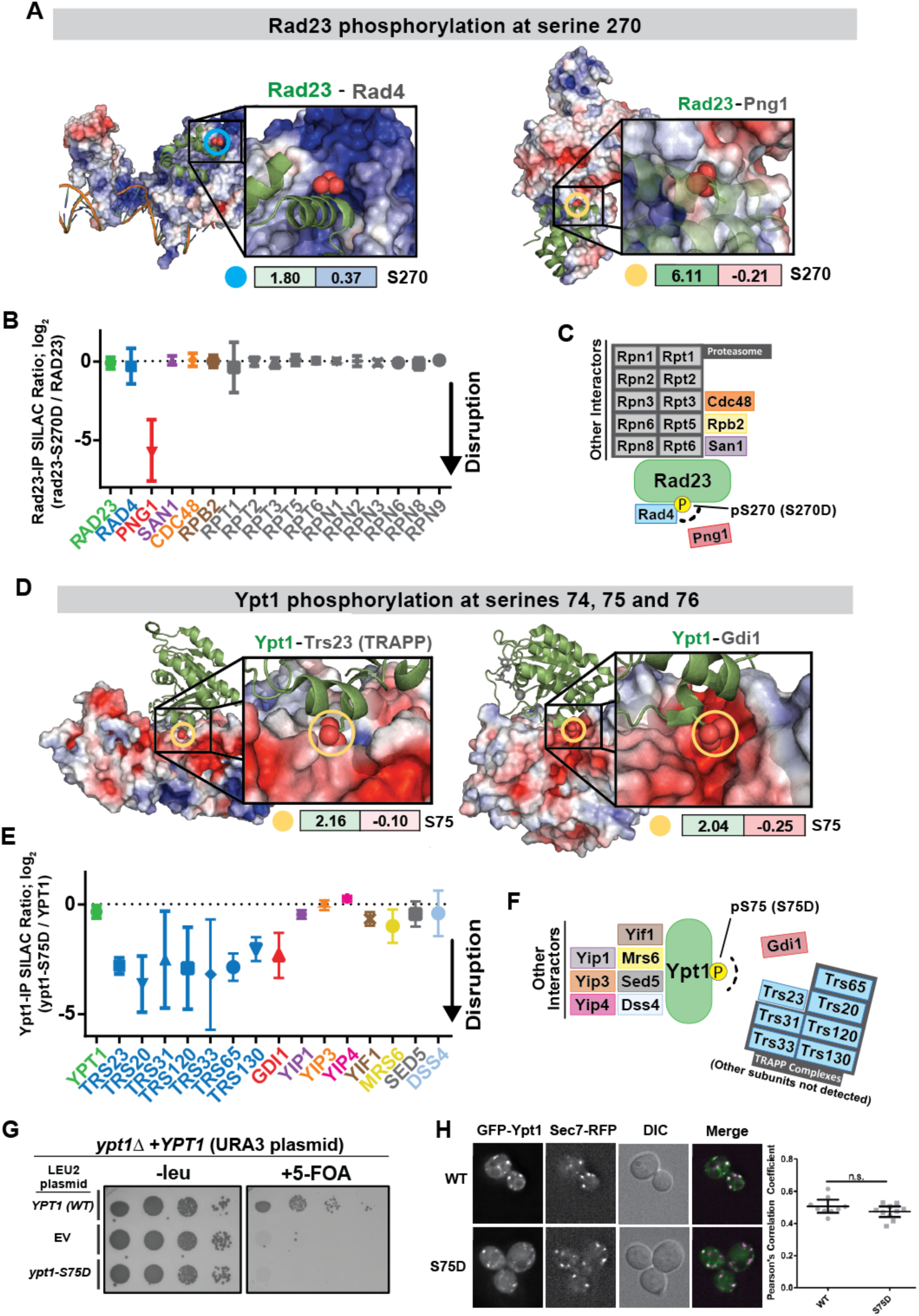
Validation of the predicted effects of phosphorylation of Rad23 and Ypt1 in disrupting specific protein-protein interactions. (A) Rad23 in complex with Rad4 (left) and Png1 (right). S (left) and E (right) scores are displayed for the phosphorylation of Rad23 at serine 270 (circled in in both structures). (B) Quantitative mass spectrometry analysis of the Rad23 interaction network and the effect of a phospho-mimetic mutation at serine 270. SILAC labeled yeast cultures expressing Rad23-FLAG or an empty vector were subjected to anit-FLAG IP to pre-define the list of specific Rad23 interacting proteins shown in the graph. The average SILAC ratios represent quantitative analysis of fold changes for each of the interactions in IP using wild-type Rad23 versus Rad23-S270D mutant as bait (Supplemental Table 8). (C) Schematics depicting Rad23’s protein interaction network and the impact of a phospho-mimetic mutation at serine 270, as defined by SILAC IP-MS. (D) Ypt1 in complex with the TRAPP complex (left) and Gdi1 (right). We detected phosphorylation at S74, S75, and S76 of Ypt1 (all with high confidence localization in singly phosphorylated phosphopeptides). S (left) and E (right) scores for the monophosphorylation Ypt1 at serine 75 is displayed and modeled into both structures (circled). (E) Quantitative mass spectrometry analysis of the Ypt1 interaction network and the effect of a phospho-mimetic mutation at serine 75. SILAC labeled yeast cultures expressing Ypt1-FLAG or an empty vector were subjected to anit-FLAG IP to pre-define the list of specific Ypt1-interacting proteins shown in the graph. The average SILAC ratios represent quantitative analysis of fold changes for each of the interactions in IP using wild-type Ypt1 versus Ypt1-S75D mutant as bait (Supplemental Table 9). (F) Schematics depicting Ypt1’s protein interaction network and the impact of a phospho-mimetic mutation at serine 75, as defined by SILAC IP-MS. (G) Plasmid shuffling assay to test the whether ypt1-S75D can fulfill Ypt1’s essential functions. (H) Fluorescent microscopy to determine whether ypt1-S75D retains proper localization to the Golgi complex. Plasmids expressing either wild-type GFP-Ypt1 or GFP-Ypt1-S75D were transformed into yeast cells. Single focal planes are shown of live-cell fluorescence microscopy images under normal growth conditions. Cells are expressing an endogenously tagged Golgi marker, Sec7-DsRed. Scale bar is 2um. The amount of overlap between Ypt1 and Sec7 colocalization was quantified using the Pearson’s Correlation Coefficient.

We performed a similar analysis for the Golgi-resident Rab family GTPase, Ypt1, the essential Rab1 homolog that regulates ER-to-Golgi membrane trafficking by recruiting effectors to the membrane surfaces of ER-derived vesicles and the Golgi complex ^33, 34^. The primary guanine-nucleotide exchange factor (GEF) that activates Ypt1 *in vivo* is the TRAPPIII complex ^35, 36^; the related TRAPPII complex is also capable of promoting Ypt1 activation ^37^. Ypt1 interacts with the TRAPP complexes, in part, by binding to the Trs23 subunit. The inactive (GDP-bound) form of Ypt1 is kept soluble in the cytoplasm by binding to Gdi1 ^38^. Gdi1 therefore prevents inactive Ypt1 from accumulating on the membrane of the Golgi complex or other organelles.

Interestingly, we found three consecutive phosphosites in Ypt1 (S74, S75, and S76) that lie at its interface with both Trs23 and Gdi1, potentially disrupting these interactions when phosphorylated (Fig. 5d). A phosphomimetic mutation of just one of these residues, S75D, was enough to disrupt the interactions of Ypt1 with subunits of the TRAPP complexes (including Trs23) and with Gdi1, without disrupting other interactions (Figs. 5e, f). We also tested whether the S75D mutation retained the essential function of Ypt1. We observed that *ypt1-S75D* was unable to support viability in the absence of endogenous Ypt1 (Fig. 5g). However, the GFP-Ypt1-S75D mutant protein appeared to retain normal Golgi localization, measured by co-localization with the Golgi-resident protein Sec7 (Fig. 5h), suggesting that S75D is a separation-of-function mutation that will be useful to dissect distinct mechanisms of Ypt1 regulation. Overall, these examples reinforce the concept that it is possible to systematically predict the impact of phosphorylation on the regulation of protein-protein interactions based on the structural context of its occurrence.

## Discussion

Here we conducted the most in depth analysis of the phosphoproteome in budding yeast to date, and our efforts have nearly doubled the number of identified phosphorylation sites in this organism. In all, if considering both our current analysis and previous reports, the budding yeast phosphoproteome presently consists of approximately 40,000 identified phosphorylation sites. While the plot in Figure 2d indicates that our ability to identify novel phosphorylation sites in the yeast phosphoproteome may be approaching its limit, we acknowledge that this saturation analysis is biased toward our instrumentation and methodologies. Nonetheless, our coverage is, in some cases, approaching the coverage achievable through the analysis of purified protein complexes (Fig. 2e), which suggests that the majority of potentially detectable phosphorylation events in budding yeast have likely been identified.

The biological significance of nearly all of these identified phosphorylation events remains unknown. The extensive scope of the phosphoproteome raises the question as to whether many of the identified phosphorylation events actually have tangible biological significance. While quantity of phosphosites identified and phosphoproteome coverage achieved in our study has inherent value, the ability to distinguish functional phosphorylation from what could potentially be “off-target” or promiscuous kinase action represents the primary challenge in dealing with large-scale phosphoproteomic datasets. Importantly, while the “detectability” of a particular phosphorylation event is impacted by factors other than its abundance (e.g. peptide solubility or ionization, accessibility to tryptic digestion, etc.), an argument could be made that many of the phosphorylation events that are buried deep within the phosphoproteome are low abundant and, therefore, are likely less important than the events which are readily detectable. If true, by expanding the depth with which the phosphoproteome is profiled, to what extent is our dataset actually revealing functional phosphorylation sites? While this question is difficult to address, the examples presented in Figure 2f strongly indicate that even phosphorylation that exists near the threshold of MS detection can be biologically meaningful. In addition, the relative prevalence of a particular phosphorylation event does not predetermine its importance. Although specialized MS applications can assess the stoichiometry of protein phosphorylation ^39^, general MS-based phosphoproteomics does not inherently inform phosphorylation stoichiometry. Therefore, low-abundant phosphoproteins that are stoichiometrically phosphorylated are often indistinguishable from very abundant proteins whose phosphorylated form represents only a small fraction of their total protein.

Given the challenges highlighted above, how can one distinguish functional biologically meaningful phosphorylation from what might just be the “noise” of the phosphoproteome? For one, it is clear that the evolutionary conservation of a phosphorylated residue does not dictate the relevancy of the event, as many functional phosphorylation events occur on poorly conserved residues ^17, 40^. One clever way to distinguish a “deliberate” phosphorylation event from promiscuous ones may be to measure how dynamically it changes. With the assumption that functional kinase-substrate interactions are better optimized for binding than promiscuous interactions, Kanshin et al. recently demonstrated that changes in phosphorylation occur faster on functional versus promiscuous substrates ^23^. Nevertheless, the current standard for determining the functionality of a phosphorylation event requires the generation of mutant yeast strains that either lack or constitutively mimic the phosphorylated residues in a substrate protein, with the ultimate goal of phenocopying the effects of a kinase’s action or inaction. However, generating phospho-site mutations is labor intensive and often times, due to the recessive nature of phospho-mutant phenotypes, requires genetic manipulation at the endogenous locus. Efforts to elicit phenotypes from phospho-site mutants can be further complicated by functional redundancy, which in some cases can be found in serines or threonines neighboring the identified phosphorylation event or in another substrate whose phosphorylation results in a redundant effect. The phospho-regulation of Slx4 and Sld3/Dbf4 exemplifies of the extensive redundancy that must be overcome when making phospho-site mutants ^41, 42^.

### 3D phosphoproteome analysis provides insights into function and regulatory mechanism

The challenges that hinder the interpretation of large phosphoproteomic datasets are, in some ways, similar to those faced in the field of human genomics. As mass spectrometry has exponentially expanded the catalog of phosphorylation events, genomics has similarly revealed tens of thousands of disease-associated mutations ^43, 44^. Akin to the biologically impactful phosphorylation events in our dataset, impactful mutations exist amidst many less meaningful polymorphisms ^45–47^. Recent efforts to identify the key mutations that underpin human disease phenotypes have utilized the expanding collection of protein structural information ^48–51^, with the logic being that mutations that occur at or near the interfaces where proteins interact will have a higher likelihood of impacting protein function. Building on this logic, here we streamlined the identification of the functional phosphosites by identifying those located at or near protein-interaction interfaces (Fig. 4). In addition, we were able to make systematic predictions about the impact of a phosphorylation event occurring near protein-protein interfaces, and its potential for causing steric clashes or an electrostatic environment incompatible with the crystal structure. Our minimalistic approach to predicting regulatory phosphorylation based on available structural information, though simpler than methods employed previously ^52–54^, was successful at predicting disruptive phosphorylation (Fig. 5). For example, we demonstrated that phosphorylation-mimicking mutation of Rad23 at S270 specifically disrupts the interaction between Rad23 and Png1. The formation of the Rad23-Png1 complex has been shown to be critical for the efficient degradation of glycosylated ER-associated proteins ^32^. Thus, while the kinase responsible for the phosphorylation of S270 and the biological context in which this phosphorylation occurs remain unclear, the phosphorylation of Rad23 at S270 could, in theory, act as a switch to inhibit the degradation of glycosylated ER proteins. Moreover, we found that phosphomimetic mutation of S75 in the Rab GTPase Ypt1 disrupts its interactions with both Gdi1 and the TRAPP GEF complexes. This suggests that phosphorylation of S75 would result in decreased activation of Ypt1 but persistence of inactive Ypt1 on the Golgi membrane. Correspondingly, the Ypt1 phosphomutant retained its localization to the Golgi yet was unable to provide the essential function of Ypt1. A previous study demonstrated that phosphorylation of another Rab GTPase, Sec4, is a negative regulatory mechanism coupled to the cell cycle ^55^. Therefore, although it remains to be determined how phosphorylation of Ypt1 is regulated, our results suggest that phosphorylation of Ypt1 is a plausible regulatory mechanism for controlling when and where it is activated.

Moving forward, the ever expanding repository of structural information will benefit methodologies that utilize that information to predict the functionality of post translational modifications. Elucidation of more protein structures, particularly through the use of emerging technologies like Cyro-EM, will expand the 3D characterization of the phosphoproteome. Thus, methods like those presented here will continue to drive the functional exploration of the phosphoproteome.

## Materials and Methods

### Protein extraction and sample preparation for phosphoproteome analysis

The phosphoproteomic experiments used as the source for the database were performed for a variety of focused biological investigations. In almost all cases these experiments were performed with the intention of quantifying changes in phosphopeptide abundance, and thus relied on a two-channel SILAC-based workflow. “Light” and “heavy”-labeled cultures (“light” version complemented with normal arginine and lysine; “heavy” version complemented with L-Lysine 13C6, 15N2.HCl and L-Arginine 13C6, 15N4.HCl) were combined, harvested by centrifugation in TE buffer pH 8.0 containing protease inhibitors and stored frozen at -80°C until cell lysis. Approximately 0.3 g of yeast cell pellet (in 3 separate 2mL screwcap tubes) was lysed by bead beating at 4°C in 3 mL of lysis buffer (1mL per tube) containing 50 mM Tris-HCl, pH 8.0, 0.2% Tergitol, 150 mM NaCl, 5 mM EDTA, complete EDTA-free protease inhibitor cocktail (Roche), 5 mM sodium fluoride and 10 mM β-glycerophosphate. Lysates of light and heavy conditions were mixed together (approximately 6mgs of protein from each condition). The mixed lysate was then denatured in 1% SDS, reduced with DTT, alkylated with iodoacetamide and then precipitated with three volumes of a solution containing 50% acetone and 50% ethanol. Proteins were solubilized in a solution of 2 M urea, 50 mM Tris-HCl, pH 8.0, and 150 mM NaCl, and then TPCK-treated trypsin was added. Digestion was performed overnight at 37°C, and then trifluoroacetic acid and formic acid were added to a final concentration of 0.2%. Peptides were desalted with Sep-Pak C18 column (Waters). C18 column was conditioned with 5 column volumes of 80% acetonitrile and 0.1% acetic acid and washed with 5 column volumes of 0.1% trifluoroacetic acid. After samples were loaded, column was washed with 5 column volumes of 0.1% acetic acid followed by elution with 4 column volumes of 80% acetonitrile and 0.1% acetic acid. Elution was dried in a SpeedVac evaporator and resuspended in 1% acetic acid.

### Phosphopeptide enrichment

After protein extraction and trypsin digestion, desalted peptides were resuspended in 1% acetic acid and loaded in a tip column containing ∼22µl of immobilized metal affinity chromatography (IMAC) resin prepared as previously described ^56^. After loading, the IMAC resin was washed with 1 column volume of 25% acetonitrile, 100 mM NaCl, and 0.1% acetic acid solution followed by 2 column volumes of 1% acetic acid, 1 column volume of deionized water and finally, eluted with 3 column volumes of 12% ammonia and 10% acetonitrile solution. The elutions were then dried and resuspended in 16.5ul H20.

### HILIC fractionation

After phosphopeptide enrichment, samples were dried in a SpeedVac, reconstituted in 80% acetonitrile and 1% formic acid and fractionated by hydrophilic interaction liquid chromatography (HILIC) with TSK gel Amide-80 column (2 mm x 150 mm, 5 µm; Tosoh Bioscience). 90sec fractions were collected between 10 and 25 min of the gradient. Three solvents were used for the gradient: buffer A (90% acetonitrile); buffer B (80% acetonitrile and 0.005% trifluoroacetic acid) and buffer C (0.025% trifluoroacetic acid). The gradient used consists of a 100% buffer A at time = 0 min; 88% of buffer B and 12% of buffer C at time = 5 min; 60% of buffer B and 40% of buffer C at time = 30 min; and 5% of buffer B and 95 % of buffer C from time = 35 to 45 min in a flow of 150 µl/min.

### Mass spectrometry analysis and data acquisition

HILIC fractions were dried in a SpeedVac, reconstituted in 0.1% trifluoroacetic acid and subjected to LC-MS/MS analysis using a 20-cm-long 125-µm inner diameter column packed in-house with 3 µm C18 reversed-phase particles (Magic C18 AQ beads, Bruker). Separated phosphopeptides were electrosprayed into a QExactive Orbitrap mass spectrometer (Thermo Fisher Scientific). Xcalibur software (Thermo Fischer Scientific) was used for the data acquisition and the Q Exactive was operated in data-dependent mode. Survey scans were acquired in the Orbitrap mass analyzer over the range of 380 to 1800 m/z with a mass resolution of 70,000 (at m/z 200). MS/MS spectra was performed selecting up to the 10 most abundant ions with a charge state using of 2, 3 or 4 within an isolation window of 2.0 m/z. Selected ions were fragmented by Higher-energy Collisional Dissociation (HCD) with normalized collision energies of 27 and the tandem mass spectra was acquired in the Orbitrap mass analyzer with a mass resolution of 17,500 (at m/z 200). Repeated sequencing of peptides was kept to a minimum by dynamic exclusion of the sequenced peptides for 30 seconds. For MS/MS, AGC target was set to 1e5 and max injection time was set to 120ms.

### Phosphopeptide and phosphorylation site identification: primary search using Andromeda

Three separate search engines were used to search the raw MS/MS spectra. All searches were performed on 19 separate “chunks”, with each chunk containing an average of 500,000 MS/MS spectra. The primary search engine used was Andromeda, as part of the MaxQuant software package (version 1.6.5.0). Searching parameters for MaxQuant included a fully-tryptic requirement. After a “first search” at 20ppm, the precursor match tolerance was set to 4.5ppm. Differential modifications were 8.0142 daltons for lysine, 10.00827 daltons for arginine, 79.966331 daltons for phosphorylation of serine, threonine and tyrosine, phosphorylation dehydration, and a static mass modification of 57.021465 daltons for alkylated cysteine residues. N-terminal acetylation was also set as a variable modification, but only for peptides that correspond to the N-terminus of protein. A complete list of searching parameters can be found in Supplemental Table 10. The primary source for the phosphosite identification was the “Phospho STY” output table in MQ ^2^. The quality threshold for a PSM to be considered for phosphosite identification was an Andromeda score greater than 40 and a delta score of greater than 6 (similar to criteria used by ^2^). The 19 “Phospho STY” files (from each of the 19 chunks) were concatenated and redundancy eliminated by retaining the PSM entry with best phosphosite localization score for every identified phosphosite. The primary dataset contains only phosphosites with high-confidence localization, which we considered as having a MaxQuant localization score of greater than 0.70.

### Phosphopeptide and phosphorylation identification: secondary search using Sequest

All spectra were also searched using two Sequest-based engines, Proteome Discoverer (Thermo) and SORCERER (Sage N Research, Inc.). For PD and SORCERER we used similar search parameters as MaxQuant, with the exception that we permitted semi-tryptic digestion, rather than require fully typtic. Precursor match tolerance for both Sequest searches was set to 10ppm. We considered only high confidence PSMs for the pipeline, filtered to less than 1% FDR using percolator and sorcererscore for PD and SORCERER, respectively. To increase the confidence in our Sequest searches further, we only considered phosphopeptides whose backbone sequence appeared in both the PD and SORCERER PSM searches. For phosphopeptides that passed this backbone requirement, we then retained the PSM information acquired using Proteome Discoverer. The phosphorylation localization probabilities were determined using ptmRS (PhosphoRS) within Proteome Discoverer ^57^. The threshold for “high-confidence” phosphosite localization was a phosphoRS percentage of >70%.

### Inclusion of high-confidence phosphosite clusters

Phosphopeptides that did not contain phosphosite localization scores that met our “high-confidence” threshold (MQ: localization score above 0.75; PD/SORCERER: phosphoRS percentage above 70%) were subsequently searched for phosphosite “clusters”. This involved identifying consecutive S/T/Y residues that were assigned localization scores which sum to greater than 0.9 (MQ) or 90% (phosphoRS) (see Supplemental Figure 1 for hypothetical examples).

### Analysis of cell-cycle phosphorylation dynamics

The spectral counts used to perform the cell cycle analysis were extracted from the primary search outlined in Figure 1. A curated set of runs were given either a G1, S phase, or G2M annotation. For every phosphosite (specifically, its best corresponding phosphopeptide), the number of identifications (i.e. spectral counts, PSMs) within runs with cell cycle annotation was tallied. Only phosphopeptides that were detected more than 5 times in the annotated runs (G1+G2M+MMS) were retained. The stringency for enrichment or depletion in a particular cell cycles state was a 5-fold difference in the number of identifications for one cell cycle stage VS the other two.

G1 synchrony was achieved through alpha factor arrest. S-phase synchrony was primarily achieved via two hour MS treatment (in some cases, 40min release from alpha factor arrest). G2/M synchrony was achieved via 2.5 hours of nocodozole treatment. The use of DNA damaging agents to synchronize cells in S phase was counterbalanced by the inclusion of multiple experiments involving the addition of 4NQO to both G1- and G2/M-arrested cell. Thus, each of the cell cycle stage contains analyses done with and without the presence of a DNA damage.

### Calculation of E and S scores for phosphosites near protein-protein interaction interfaces

All the phosphosites were mapped onto the available 3D structures in PDB ^58^ using residue-level mapping information obtained from SIFTS database ^59^. The sites that were mapped to structures with more than one chain were further considered for calculation of E and S scores. For each site, neighboring atoms with in a distance of 10Å were identified (excluding the atoms present in the same chain as the phosphosite) using an in-house python script (Atoms chosen to be the “phosphosite atoms”: “OG” for serine, “OG1” for threonine and “OH” for tyrosine). Next, the charge for all the individual neighboring atoms was obtained using a command line version of PDB2PQR pipeline (with amber as the forcefield and using ‘--nodebump’ option). Finally, the S and E scores were calculated using the following equations:

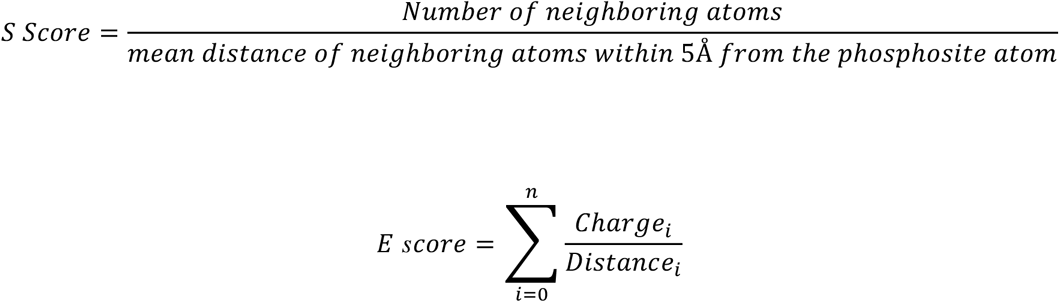

Where, *Charge_i_* is the charge of neighboring atom *i* and *Distance_i_* is the distance between atom *i* and the phosphosite atom.

### Quantitative MS analysis pull-down protein complexes

Yeast carrying either GFP-YPT1 or Rad23-FLAG were grown to an O.D.600 of 0.4 in 200 mL of -Arg -Lys dropout media (“light” version complemented with normal arginine and lysine; “heavy” version complemented with L-Lysine 13C6, 15N2.HCl and L-Arginine 13C6, 15N4.HCl). After centrifugation, pellets were kept at -80C prior to cell lysis. Approximately 0.3 g of cell pellet of each strain was lysed by bead beating at 4°C in 3 mL of lysis buffer (50 mM Tris-HCl pH 7.5, 0.2% Tergitol, 150 mM NaCl, 5 mM EDTA, Complete EDTA-free protease inhibitor cocktail (Roche), 5 mM sodium fluoride, 10 mM B-glycerol-phosphate). Lysates were incubated with GFP-TRAP (in-house) or anti-FLAG agarose resin (Sigma) for 4 hours at 4°C. After 3 washes with lysis buffer, bound proteins were eluted with 90uls of elution buffer (100 mM Tris-HCl pH 8.0, 1% SDS). Eluted proteins from normal or heavy media grown cells were mixed together, reduced, alkylated and then precipitated with three volumes of a solution containing 50% acetone and 50% ethanol. Proteins were solubilized in a solution of 2 M urea, 50 mM Tris-HCl, pH 8.0, and 150 mM NaCl, and then Trypsin Gold was added. Digestion was performed overnight at 37°C, and then trifluoroacetic acid and formic acid were added to a final concentration of 0.2%. Peptides were desalted with Sep-Pak C18 column (Waters). Elution from C18 column was dried in a SpeedVac evaporator and resuspended in 0.1% trifluoroacetic acid.

### Fluorescent Microscopy

Overnight cultures were grown to an OD600 between 0.1 and 0.8, then imaged. Single focal planes are shown of live-cell fluorescence microscopy images under normal growth conditions. Cells are expressing endogenously tagged Sec7-6xDsRed. Scale bar is 2um. The amount of overlap between Ypt1 and Sec7 was quantified using the Pearson’s Correlation Coefficient. A region of interest was selected surrounding 1-4 cells and propagated to all of the focal planes containing those cells for correlation analysis. Each data point represents the PCC for an image (WT = 9 images, S75D = 10 images) containing several regions of interest totaling 4-21 cells. A total of 105 cells for WT and S75D, while 107 cells for S39D were analyzed. An unpaired two-tailed t-test with Welch’s correction was used to analyze the data points. The PCC for the WT and S75D mutant are not significantly different (p=0.1667), but S39D is significantly different from both WT and S75D (p < 0.0001, and p = 0.0002). WT Mean = 0.5068, S75D Mean = 0.4732, S39D Mean = 0.3772, error bars represent 95% CIs.

## Acknowledgements

We would like to thank Bik Tye and Yuanliang Zhai for critical reading of the manuscript. Mimi Xie assisted in making the ternary plot. Kiran Madura graciously provided Rad23-related plasmids. Adm Chrysler provided invaluable IT assistance. Finally, we thank members of the Smolka and Yu Labs for useful discussions and constructive comments.

This work was supported by grants from the National Institutes of Health to M.B.S. (R01GM097272 and R01HD095296; Equipment Supplement R01GM097272-07S1) and H.Y. (R01GM124559 and R01GM125639).

## Author contributions

M.C.L. performed all phosphoproteomic experiments. M.C.L., E.J.S., and S. V. performed IP-MS experiments. M.C.L, K.Y., M.B.S., V.T., H.Y. conceived the data processing pipeline. K.Y. and S.G. wrote various scripts for the consolidation and interpretation of the dataset and also designed the website. K.Y. wrote the clash scoring algorithm. M.C.L. and M.B.S. wrote the manuscript.

## References

1. Aebersold, R. & Mann, M. Mass spectrometry-based proteomics. Nature 422, 198–207 (2003).

2. Sharma, K. et al. Ultradeep human phosphoproteome reveals a distinct regulatory nature of Tyr and Ser/Thr-based signaling. Cell Rep 8, 1583–94 (2014).

3. Olsen, J.V. et al. Global, in vivo, and site-specific phosphorylation dynamics in signaling networks. Cell 127, 635–48 (2006).

4. Swaney, D.L. et al. Global analysis of phosphorylation and ubiquitylation cross-talk in protein degradation. Nat Methods 10, 676–82 (2013).

5. Bastos de Oliveira, F.M., et al. Phosphoproteomics reveals distinct modes of Mec1/ATR signaling during DNA replication. Mol Cell 57, 1124–32 (2015).

6. Needham, E.J., Parker, B.L., Burykin, T., James, D.E. & Humphrey, S.J. Illuminating the dark phosphoproteome. Sci Signal 12(2019).

7. Hornbeck, P.V. et al. 15 years of PhosphoSitePlus(R): integrating post-translationally modified sites, disease variants and isoforms. Nucleic Acids Res 47, D433–D441 (2019).

8. Lienhard, G.E. Non-functional phosphorylations? Trends Biochem Sci 33, 351–2 (2008).

9. Landry, C.R., Levy, E.D. & Michnick, S.W. Weak functional constraints on phosphoproteomes. Trends Genet 25, 193–7 (2009).

10. Dephoure, N., Gould, K.L., Gygi, S.P. & Kellogg, D.R. Mapping and analysis of phosphorylation sites: a quick guide for cell biologists. Mol Biol Cell 24, 535–42 (2013).

11. Levy, E.D., Michnick, S.W. & Landry, C.R. Protein abundance is key to distinguish promiscuous from functional phosphorylation based on evolutionary information. Philos Trans R Soc Lond B Biol Sci 367, 2594–606 (2012).

12. Lanz, M.C. et al. Separable roles for Mec1/ATR in genome maintenance, DNA replication, and checkpoint signaling. Genes Dev 32, 822–835 (2018).

13. Thompson, A.J., Abu, M. & Hanger, D.P. Key issues in the acquisition and analysis of qualitative and quantitative mass spectrometry data for peptide-centric proteomic experiments. Amino Acids 43, 1075–85 (2012).

14. Oughtred, R. et al. The BioGRID interaction database: 2019 update. Nucleic Acids Res 47, D529–D541 (2019).

15. Ho, B., Baryshnikova, A. & Brown, G.W. Unification of Protein Abundance Datasets Yields a Quantitative Saccharomyces cerevisiae Proteome. Cell Syst 6, 192–205 e3 (2018).

16. Solari, F.A., Dell’Aica, M., Sickmann, A. & Zahedi, R.P. Why phosphoproteomics is still a challenge. Mol Biosyst 11, 1487–93 (2015).

17. Amoutzias, G.D., He, Y., Lilley, K.S., Van de Peer, Y. & Oliver, S.G. Evaluation and properties of the budding yeast phosphoproteome. Mol Cell Proteomics 11, M111 009555 (2012).

18. Albuquerque, C.P. et al. A multidimensional chromatography technology for in-depth phosphoproteome analysis. Mol Cell Proteomics 7, 1389–96 (2008).

19. Blanco, M.G., Matos, J. & West, S.C. Dual control of Yen1 nuclease activity and cellular localization by Cdk and Cdc14 prevents genome instability. Mol Cell 54, 94–106 (2014).

20. Rossi, S.E., Ajazi, A., Carotenuto, W., Foiani, M. & Giannattasio, M. Rad53-Mediated Regulation of Rrm3 and Pif1 DNA Helicases Contributes to Prevention of Aberrant Fork Transitions under Replication Stress. Cell Rep 13, 80–92 (2015).

21. Kono, K. et al. G1/S cyclin-dependent kinase regulates small GTPase Rho1p through phosphorylation of RhoGEF Tus1p in Saccharomyces cerevisiae. Mol Biol Cell 19, 1763–71 (2008).

22. Li, Z. et al. Systematic exploration of essential yeast gene function with temperature-sensitive mutants. Nat Biotechnol 29, 361–7 (2011).

23. Kanshin, E., Bergeron-Sandoval, L.P., Isik, S.S., Thibault, P. & Michnick, S.W. A cell-signaling network temporally resolves specific versus promiscuous phosphorylation. Cell Rep 10, 1202–14 (2015).

24. Swaffer, M.P., Jones, A.W., Flynn, H.R., Snijders, A.P. & Nurse, P. Quantitative Phosphoproteomics Reveals the Signaling Dynamics of Cell-Cycle Kinases in the Fission Yeast Schizosaccharomyces pombe. Cell Rep 24, 503–514 (2018).

25. Touati, S.A., Kataria, M., Jones, A.W., Snijders, A.P. & Uhlmann, F. Phosphoproteome dynamics during mitotic exit in budding yeast. EMBO J 37(2018).

26. Meyer, M.J. et al. Interactome INSIDER: a structural interactome browser for genomic studies. Nat Methods 15, 107–114 (2018).

27. Mosca, R., Ceol, A. & Aloy, P. Interactome3D: adding structural details to protein networks. Nat Methods 10, 47–53 (2013).

28. Iakoucheva, L.M. et al. The importance of intrinsic disorder for protein phosphorylation. Nucleic Acids Res 32, 1037–49 (2004).

29. Dolinsky, T.J., Nielsen, J.E., McCammon, J.A. & Baker, N.A. PDB2PQR: an automated pipeline for the setup of Poisson-Boltzmann electrostatics calculations. Nucleic Acids Res 32, W665–7 (2004).

30. Schauber, C. et al. Rad23 links DNA repair to the ubiquitin/proteasome pathway. Nature 391, 715–8 (1998).

31. de Laat, W.L., Jaspers, N.G. & Hoeijmakers, J.H. Molecular mechanism of nucleotide excision repair. Genes Dev 13, 768–85 (1999).

32. Kim, I. et al. The Png1-Rad23 complex regulates glycoprotein turnover. J Cell Biol 172, 211–9 (2006).

33. Hutagalung, A.H. & Novick, P.J. Role of Rab GTPases in membrane traffic and cell physiology. Physiol Rev 91, 119–49 (2011).

34. Yang, X.Z. et al. Rab1 in cell signaling, cancer and other diseases. Oncogene 35, 5699–5704 (2016).

35. Wang, W., Sacher, M. & Ferro-Novick, S. TRAPP stimulates guanine nucleotide exchange on Ypt1p. J Cell Biol 151, 289–96 (2000).

36. Thomas, L.L., Joiner, A.M.N. & Fromme, J.C. The TRAPPIII complex activates the GTPase Ypt1 (Rab1) in the secretory pathway. J Cell Biol 217, 283–298 (2018).

37. Thomas, L.L. & Fromme, J.C. GTPase cross talk regulates TRAPPII activation of Rab11 homologues during vesicle biogenesis. J Cell Biol 215, 499–513 (2016).

38. Garrett, M.D., Zahner, J.E., Cheney, C.M. & Novick, P.J. GDI1 encodes a GDP dissociation inhibitor that plays an essential role in the yeast secretory pathway. EMBO J 13, 1718–28 (1994).

39. Lim, M.Y., O’Brien, J., Paulo, J.A. & Gygi, S.P. Improved Method for Determining Absolute Phosphorylation Stoichiometry Using Bayesian Statistics and Isobaric Labeling. J Proteome Res 16, 4217–4226 (2017).

40. Holt, L.J. et al. Global analysis of Cdk1 substrate phosphorylation sites provides insights into evolution. Science 325, 1682–6 (2009).

41. Ohouo, P.Y., Bastos de Oliveira, F.M., Almeida, B.S. & Smolka, M.B. DNA damage signaling recruits the Rtt107-Slx4 scaffolds via Dpb11 to mediate replication stress response. Mol Cell 39, 300–6 (2010).

42. Zegerman, P. & Diffley, J.F. Checkpoint-dependent inhibition of DNA replication initiation by Sld3 and Dbf4 phosphorylation. Nature 467, 474–8 (2010).

43. Landrum, M.J. et al. ClinVar: public archive of relationships among sequence variation and human phenotype. Nucleic Acids Res 42, D980–5 (2014).

44. Stenson, P.D. et al. Human Gene Mutation Database (HGMD): 2003 update. Hum Mutat 21, 577–81 (2003).

45. The 1000 Genomes Project Consortium. An integrated map of genetic variation from 1,092 human genomes. Nature 491, 56–65 (2012).

46. Tennessen, J.A. et al. Evolution and functional impact of rare coding variation from deep sequencing of human exomes. Science 337, 64–69 (2012).

47. The Genome of the Netherlands Consortium. Whole-genome sequence variation, population structure and demographic history of the Dutch population. Nature Genetics 46, 818–825 (2014).

48. Wang, X. et al. Three-dimensional reconstruction of protein networks provides insight into human genetic disease. Nat Biotechnol 30, 159–64 (2012).

49. Wei, X. et al. A massively parallel pipeline to clone DNA variants and examine molecular phenotypes of human disease mutations. PLoS Genet 10, e1004819 (2014).

50. Meyer, M.J. et al. Mutation3D: Cancer Gene Prediction Through Atomic Clustering of Coding Variants in the Structural Proteome. Hum Mutat (2016).

51. Chen, S. et al. An interactome perturbation framework prioritizes damaging missense mutations for developmental disorders. Nat Genet 50, 1032–1040 (2018).

52. Betts, M.J. et al. Systematic identification of phosphorylation-mediated protein interaction switches. PLoS Comput Biol 13, e1005462 (2017).

53. Beltrao, P. et al. Systematic functional prioritization of protein posttranslational modifications. Cell 150, 413–25 (2012).

54. Nishi, H., Hashimoto, K. & Panchenko, A.R. Phosphorylation in protein-protein binding: effect on stability and function. Structure 19, 1807–15 (2011).

55. Lepore, D., Spassibojko, O., Pinto, G. & Collins, R.N. Cell cycle-dependent phosphorylation of Sec4p controls membrane deposition during cytokinesis. J Cell Biol 214, 691–703 (2016).

56. Bastos de Oliveira, F.M., Kim, D., Lanz, M. & Smolka, M.B. Quantitative Analysis of DNA Damage Signaling Responses to Chemical and Genetic Perturbations. Methods Mol Biol 1672, 645–660 (2018).

57. Taus, T. et al. Universal and confident phosphorylation site localization using phosphoRS. J Proteome Res 10, 5354–62 (2011).

58. Gilliland, G. et al. The Protein Data Bank. Nucleic Acids Research 28, 235–242 (2000).

59. Gutmanas, A. et al. SIFTS: updated Structure Integration with Function, Taxonomy and Sequences resource allows 40-fold increase in coverage of structure-based annotations for proteins. Nucleic Acids Research 47, D482–D489 (2018).

