## Supplemental Figures for "In-depth and 3-Dimensional Exploration of the Budding Yeast Phosphoproteome"

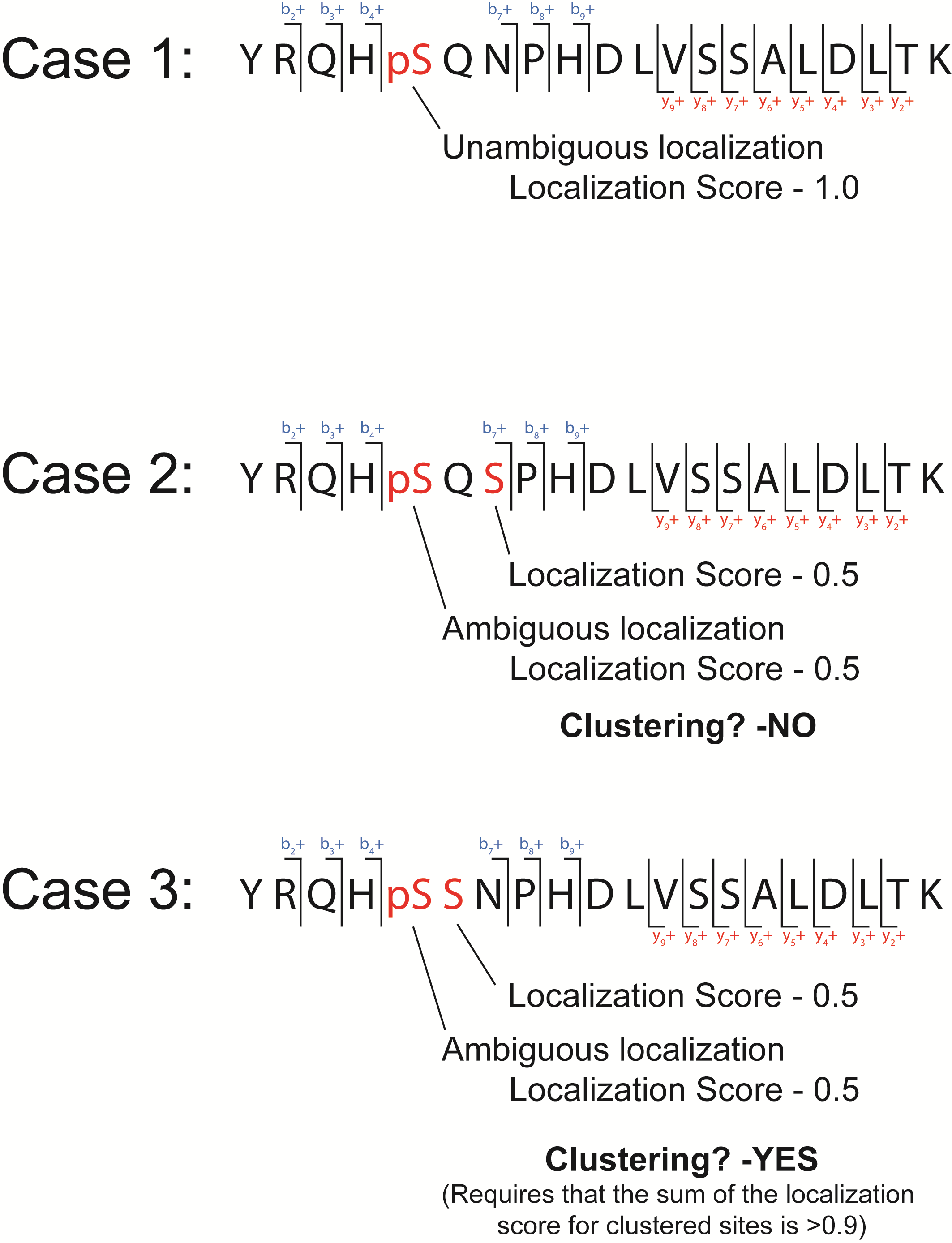

**Supplemental Figure 1 –** Schematic demonstrating the logic for the inclusion of clustered phosphosites in the dataset.

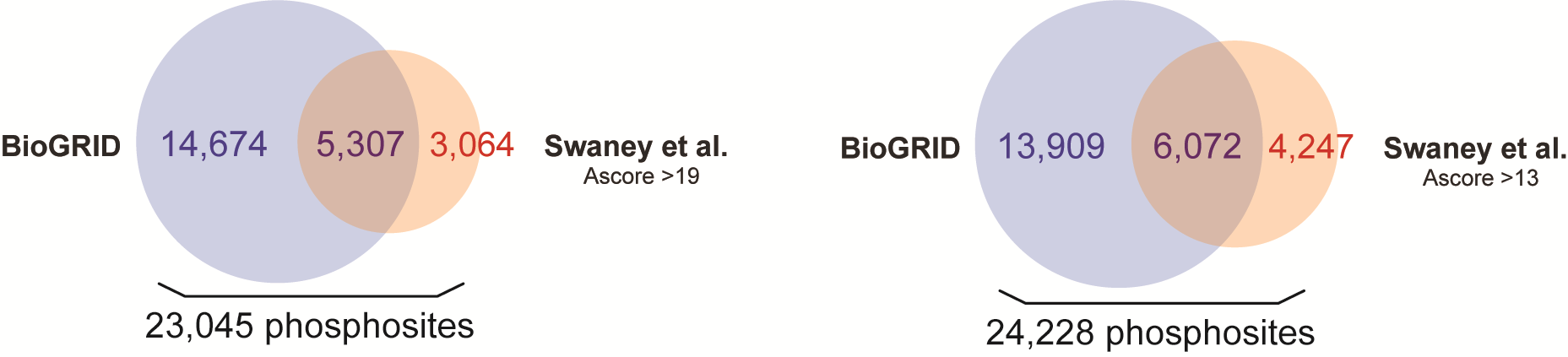

**Supplemental Figure 2 –** Compilation of the previously known phosphoproteome. For the comparison performed in Figure 2, we used the Swaney et al. data set with the more stringent cutoff for phosphosite localization.
